# Monorhinal and Birhinal Odor Processing in Humans: an fMRI investigation

**DOI:** 10.1101/2023.08.01.551475

**Authors:** Anupa Ekanayake, Qing Yang, Sangam Kanekar, Biyar Ahmed, Silas McCaslin, Deepak Kalra, Paul Eslinger, Prasanna Karunanayaka

**Author notes:** **Address correspondences to:** Prasanna R. Karunanayaka, Ph.D., Department of Radiology (Center for NMR Research) Penn State College of Medicine, The M. S. Hershey Medical Center, 500 University Dr., Hershey, PA 17033, USA.

## Abstract

The olfactory nerve, also known as cranial nerve I, is known to have exclusive ipsilateral projections to primary olfactory cortical structures. It is still unclear whether these projections also correspond to functional pathways of odor processing. In an olfactory functional magnetic resonance imaging (fMRI) study of twenty young healthy subjects with a normal sense of smell, we tested whether nostril specific stimulation with phenyl ethyl alcohol (PEA), a pure olfactory stimulant, asymmetrically activates primary or secondary olfactory-related brain structures such as primary olfactory cortex, entorhinal cortex, and orbitofrontal cortex. The results indicated that without a challenging olfactory task, passive (no sniffing) and active (with sniffing) nostril-specific PEA stimulation did not produce asymmetrical fMRI activation in olfactory cortical structures.

## INTRODUCTION

The conventional definition of the primary olfactory cortex (POC) includes brain regions that receive direct input from the olfactory bulb. Regions within the olfactory cortex have back projections to the olfactory bulb. There are strong commissural projections (via the anterior commissure) between bilateral olfactory cortical regions. Often, the POC is referred to as the piriform cortex, which is heterogeneous and located bilaterally in the frontal and temporal lobes. These frontal (anterior piriform cortex) and temporal (posterior piriform cortex) regions of the piriform are implicated in different functionality (1). Gottfried et al. postulated that the activity in the anterior piriform cortex corresponds to the perception of an odor while posterior piriform cortex activity is associated with processing the quality aspect of a chemosensory percept (2-4). Thus, the olfactory system’s anatomical connectivity pattern is able to encode a variety of complex olfactory associations (5). Although olfactory sensory neurons project exclusively to the ipsilateral olfactory bulb, cortical neurons can access bilateral input from the nasal cavity (6-8).

Compared to other species, humans have been postulated to have poor sense of smell and fewer functional olfactory receptor genes (9, 10). However, recent research contradicts this idea, stating that human olfaction is not as poor as previously thought. Studies show that humans can extract more information from their sense of smell than they consciously realize (11-13).

The anatomical morphology and functional organization of the olfactory pathway have several features that are *sui generis* among human senses. An interesting difference in the olfactory pathway is the lack of an early precortical thalamic relay to transfer peripheral input into the brain. However, some argue that the olfactory bulb plays an equivalent role to that of the thalamus since these two regions have similar structures and functions (14). While all other senses project mainly contralaterally from sensory organs into the brain, the olfactory system projects ipsilaterally. It is of interest that the spatial organization of the olfactory system is more dispersed throughout several brain regions when compared to other senses (12). Additionally, the primary cortical region in other senses typically consists of one discrete cortical area, but the POC includes a set of brain structures, some of which are subcortical (15-18).

The olfactory pathway starts with olfactory receptor cells, where volatile molecules or “odorants” activate chemoreceptors in the olfactory epithelium at the roof of the nasal cavity. These neurons pass through the lamina propria and group together into bundles called olfactory fila, which collectively make up the olfactory nerve. The olfactory nerve passes through the cribriform plate and terminates on olfactory glomeruli, which lie just beneath the surface of the bulb. The core of an olfactory glomerulus is comprised of the axons of olfactory receptor neurons, which branch and synapse to the primary dendrites of mitral and tufted cells (6). These olfactory glomeruli are also found on the olfactory bulb.

The olfactory bulb, a forebrain structure, lies along the ventral surface of the frontal lobe in the olfactory sulcus and is attached to the rest of the brain by the olfactory tract. The olfactory tract is a structure that contains fibers of the lateral olfactory tract, cells of the anterior olfactory nucleus, and fibers of the anterior limb of the anterior commissure. The latter part of the olfactory tract falls in an area where many afferent centrifugal fibers also reside and reach the olfactory bulb such as the locus coeruleus, raphe nuclei, and ipsilateral fibers from the anterior olfactory nucleus and the diagonal band.

The olfactory tract contains fibers that course caudally to terminate in areas on the telencephalon’s ventral surface, which are broadly defined as the olfactory cortex. The principal areas included in the olfactory cortex are the anterior olfactory nucleus, olfactory tubercle, piriform cortex, anterior cortical amygdaloid nucleus, periamygdaloid cortex, and lateral entorhinal cortex. These brain structures project to other structures, including the caudal orbito-frontal cortex (OFC), agranular insula, hippocampus, dorsomedial nucleus of the thalamus, medial and lateral hypothalamus, ventral striatum, and pallidum (5, 7).

*In vivo* human imaging studies (both fMRI and PET) have reported symmetric or asymmetric POC activation for birhinal stimulation or similar levels of ipsilateral and contralateral activation during monorhinal stimulation (11, 13, 19-21). Those studies, however, confounded olfactory processing with concurrent perceptual or cognitive-motor task performance, which may have biased the olfactory system laterality (19). Given the complex anatomical connectivity of the olfactory system, this study was designed to test the hypothesis that nostril-specific stimulation of a pure olfactory stimulant in trials that do not require concurrent perceptual or cognitive-motor processing will elicit bilateral activity in primary and secondary (i.e., cortical) olfactory structures regardless of sniffing.

## METHOD

### Participants

Twenty healthy normosmic volunteers (12 F), between the ages of 23-40 (mean age = 29.5 years) participated in the study. They were compensated for their travel and time. Study protocols were approved by the Penn State College of Medicine Internal Review Board (IRB) and carried out in accordance with the Declaration of Helsinki.

### Odor stimulation

Participants were MRI scanned while the odorant was delivered monorhinally (left or right nostril) or birhinally. The experiment used phenyl ethyl alcohol (PEA), which is a pure olfactory odorant, at a single concentration (1% PEA in mineral oil) (22). PEA was delivered by a computer-controlled air dilution olfactometer (ETT, Inc.) with a constant flow rate of 8 L/min. The air flow was divided between the left and right nostrils (4 L/min each nostril). The olfactometer was configured to stimulate the left or right nostril only while maintaining a fresh air flow to the opposing nostril or stimulate both nostrils simultaneously. Participants were visually cued at the start of each trial (**Figure 1**). PEA was presented for a duration of 6 s with an inter-stimulus interval was 24 s and six repetitions for each trial type. There were two runs. In the first run, subjects were instructed to inhale PEA while maintaining a normal breathing pattern. In the second run, subjects were instructed to “sniff” at the start of each PEA stimulation block. Both runs had identical imaging parameters and subjects did not perform any perceptual or cognitive-motor tasks.

**Figure 1.**
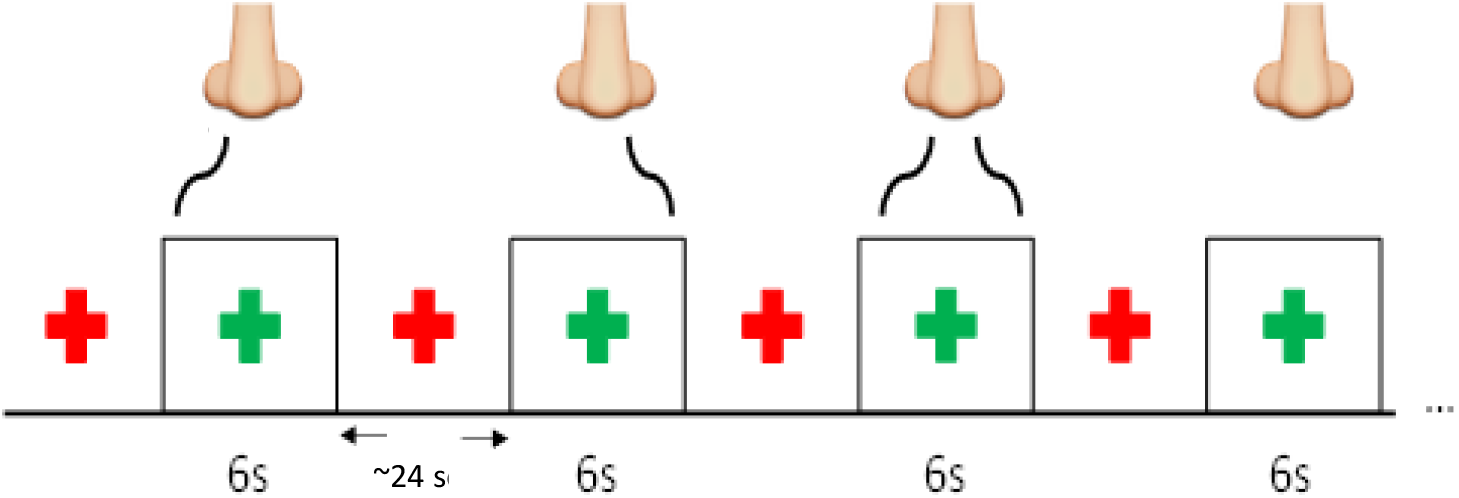
The structure of the fMRI paradigm with right only, left only, bilateral, and neither nostril stimulation with the PEA.

### MRI scanning

MRI data were obtained with a Siemens 3T Tim Trio whole-body MR scanner equipped with a 12-channel InVivo RF head coil. A T1 anatomical image was also obtained from each participant for functional overlay. fMRI data were obtained (T2*-weighted interleaved ascending GE-EPI) with following parameters: 288 volumes, 2.85 x 2.85 in plane resolution, 4 mm slice thickness with no gap, TR=2 s, TE=30 ms, FOV= 230 mm, 80 x 80 matrix size, and 90° flip angle.

### Data analysis

MRI data were analyzed with SPM8 (Wellcome Trust, London, UK). Anatomical images were segmented and transformed to Montreal Neurological Institute (MNI) standard space. Functional images were corrected for slice-timing acquisition offsets, corrected for motion, co-registered to anatomical images and transformed to MNI space, and smoothed (8 mm^3^ FWHM). Each stimulus condition was represented as a boxcar function and convolved with a hemodynamic response for first level analysis. First, we investigated activation patterns within the selected ROIs shown in **Figure 2**.

**Figure 2.**
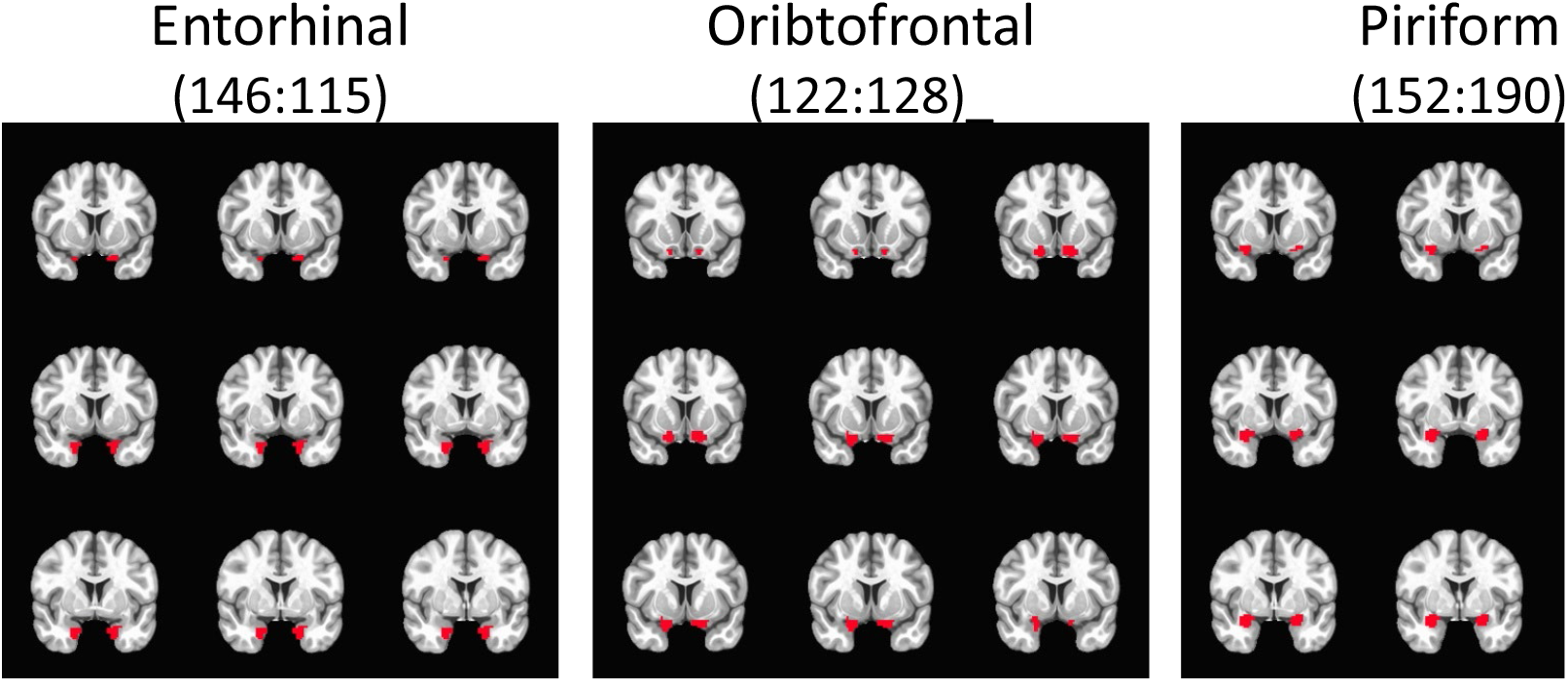
Region of interest (ROI) Masks.

As previously mentioned, the piriform cortex is heterogeneous (consisting of anterior and posterior regions) and implicated in different functionality (1). Therefore, to further investigate piriform activity patterns in our study, we performed a second level, random effects analyses with a series of t-tests using group average beta values which were family-wise error (FWE) corrected for multiple comparisons using small volume correction (SVC). In this analysis, the SVC masks included the bilateral frontal piriform, bilateral temporal piriform, and bilateral entorhinal cortex. Respective masks were created using a clustering technique on resting state fMRI (rs-fMRI) data within a larger probabilistic mask of the anatomically defined POC that was traced on a healthy human T1 image by a radiologist (**Figure 2**) (23). Functional parcellation of the POC was performed via ICA decomposition dividing the POC into three functional sub-regions bilaterally, with each region corresponding to the piriform cortex (*frontal and temporal*) and entorhinal cortex as shown in **Figure 3**. Using time courses from theses region of interests (ROIs), effective connectivity between them was estimated via extended unified structural equation modeling (euSEM) as described in Gates et al, 2011 (24).

**Figure 3.**
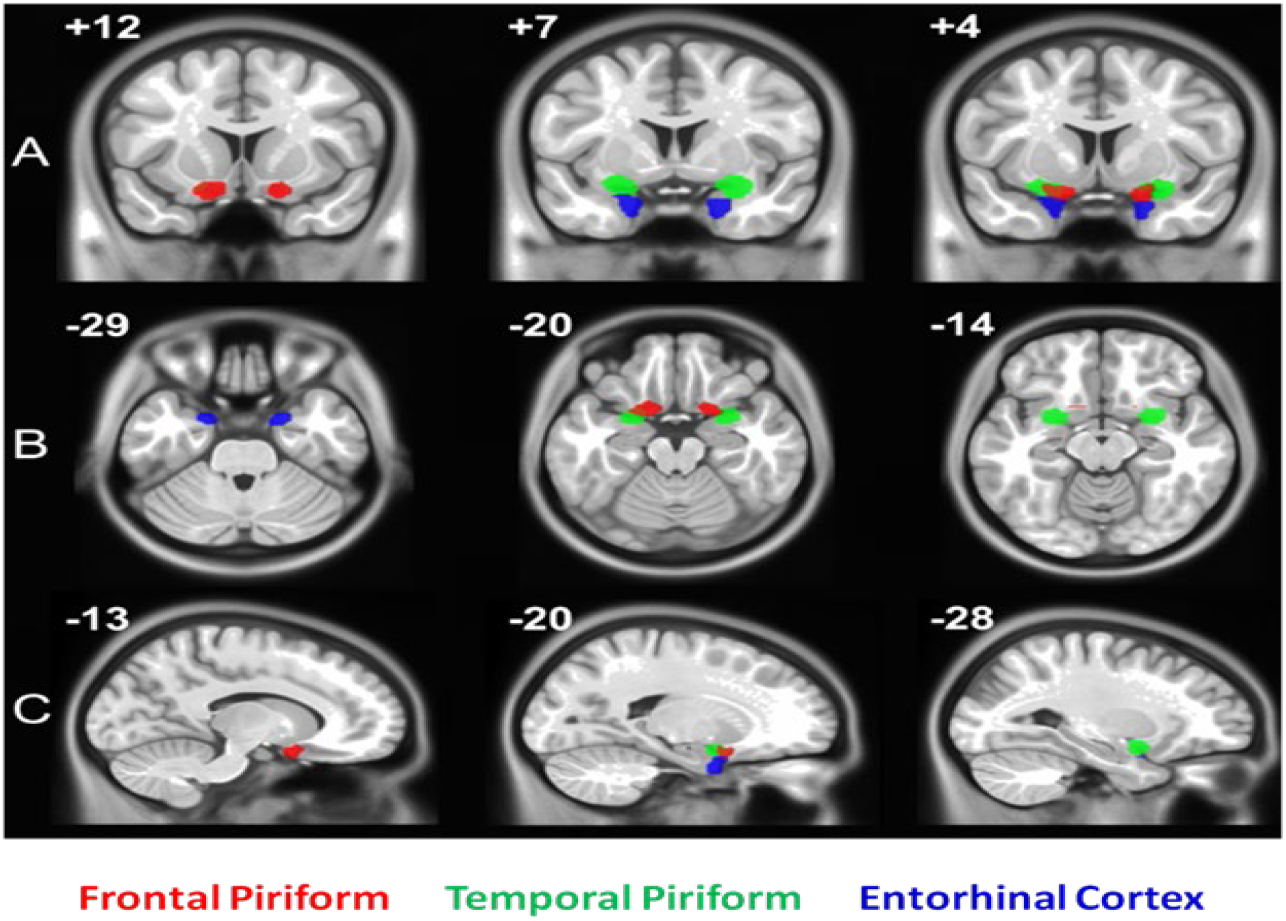
Region of interest (ROI) Masks.

## RESULTS

Figure 4 shows the group level activation maps for no-sniff and sniff paradigms and for each PEA stimulation condition. The percentage of activated voxels within ROIs are shown in **Figure 5**. The locations of peak activated voxels in each ROI that survived FWE small volume correction (SVC) to *p*< 0.05 are listed in **Table 1**.

**Table 1.** Normal breathing odorants (no-sniff paradigm)

| Nostril | Anatomy | MNI (x y z) | *t* | *z(t)* | *p(FWE)* | *k* |
| --- | --- | --- | --- | --- | --- | --- |
| Left | Left Piriform | -28 2 -20 | 4.07 | 3.86 | .002 | 91 |
|  | Right Piriform | 28 6 -22 | 3.41 | 3.28 | .012 | 149 |
|  | Left Entorhinal | -28 2 -24 | 3.91 | 3.72 | .001 | 7 |
|  | Right OFC | 24 6 -14 | 2.98 | 2.89 | .032 | 8 |
| Right | Left Piriform | -30 0 -18 | 3.29 | 3.17 | .017 | 64 |
|  | Right Piriform | 30 8 -20 | 3.69 | 3.53 | .005 | 143 |
|  | Left Entorhinal | -28 2 -24 | 2.70 | 2.63 | .03 | 2 |
|  | Right OFC | 24 8 -16 | 3.52 | 3.38 | .008 | 23 |
| Bilateral | Left Piriform | -20 0 -14 | 3.85 | 3.67 | .003 | 125 |
|  | Right Piriform | 26 8 -20 | 4.84 | 4.08 | <.001 | 184 |
|  | Left Entorhinal | -28 2 -24 | 3.23 | 3.11 | .006 | 7 |
|  | Right OFC | 24 8 -20 | 4.62 | 4.32 | <.001 | 27 |
|  | Left OFC | -24 8 -18 | 2.98 | 2.89 | .032 | 2 |

**Table 2.** Sniffing odorants (sniff paradigm)

| Nostril | Anatomy | MNI (x y z) | *t* | *z(t)* | *p(FWE)* | *k* |
| --- | --- | --- | --- | --- | --- | --- |
| Left | Left Piriform | -24 0 -18 | 3.63 | 3.48 | .005 | 69 |
|  | Right Piriform | 24 0 -18 | 3.75 | 3.58 | .004 | 87 |
|  | Left Entorhinal | -26 4 -24 | 3.23 | 3.12 | .006 | 7 |
|  | Right OFC | 24 6 -18 | 3.09 | 2.99 | .02 | 12 |
| Right | Left Piriform | -26 0 -18 | 3.32 | 3.20 | .013 | 58 |
|  | Right Piriform | 20 2 -16 | 3.70 | 3.53 | .004 | 68 |
|  | Left Entorhinal | -26 4 -24 | 3.17 | 3.07 | .008 | 7 |
|  | Right OFC | 20 6 -16 | 2.99 | 2.90 | .026 | 12 |
| Bilateral | Left Piriform | -26 4 -22 | 4.77 | 4.44 | <.001 | 97 |
|  | Right Piriform | 24 0 -20 | 4.05 | 3.84 | .001 | 85 |
|  | Left Entorhinal | -26 6 -24 | 4.98 | 4.62 | <.001 | 7 |
|  | Right OFC | 22 6 -20 | 2.88 | 2.80 | .034 | 6 |
|  | Left OFC | -22 8 -20 | 2.77 | 2.70 | .044 | 1 |

**Figure 4.**
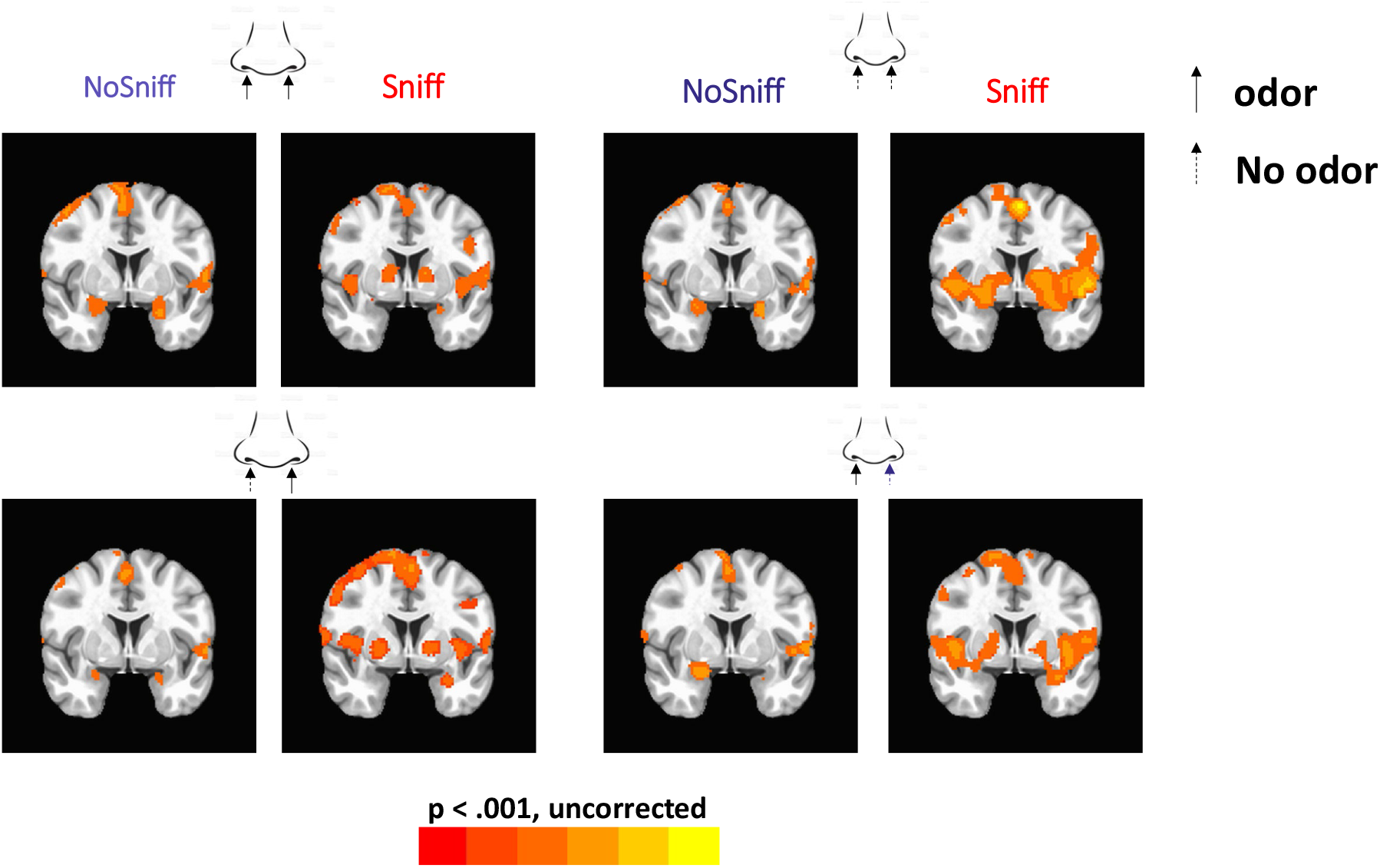
The general linear model (GLM) activation patterns.

**Figure 5.**
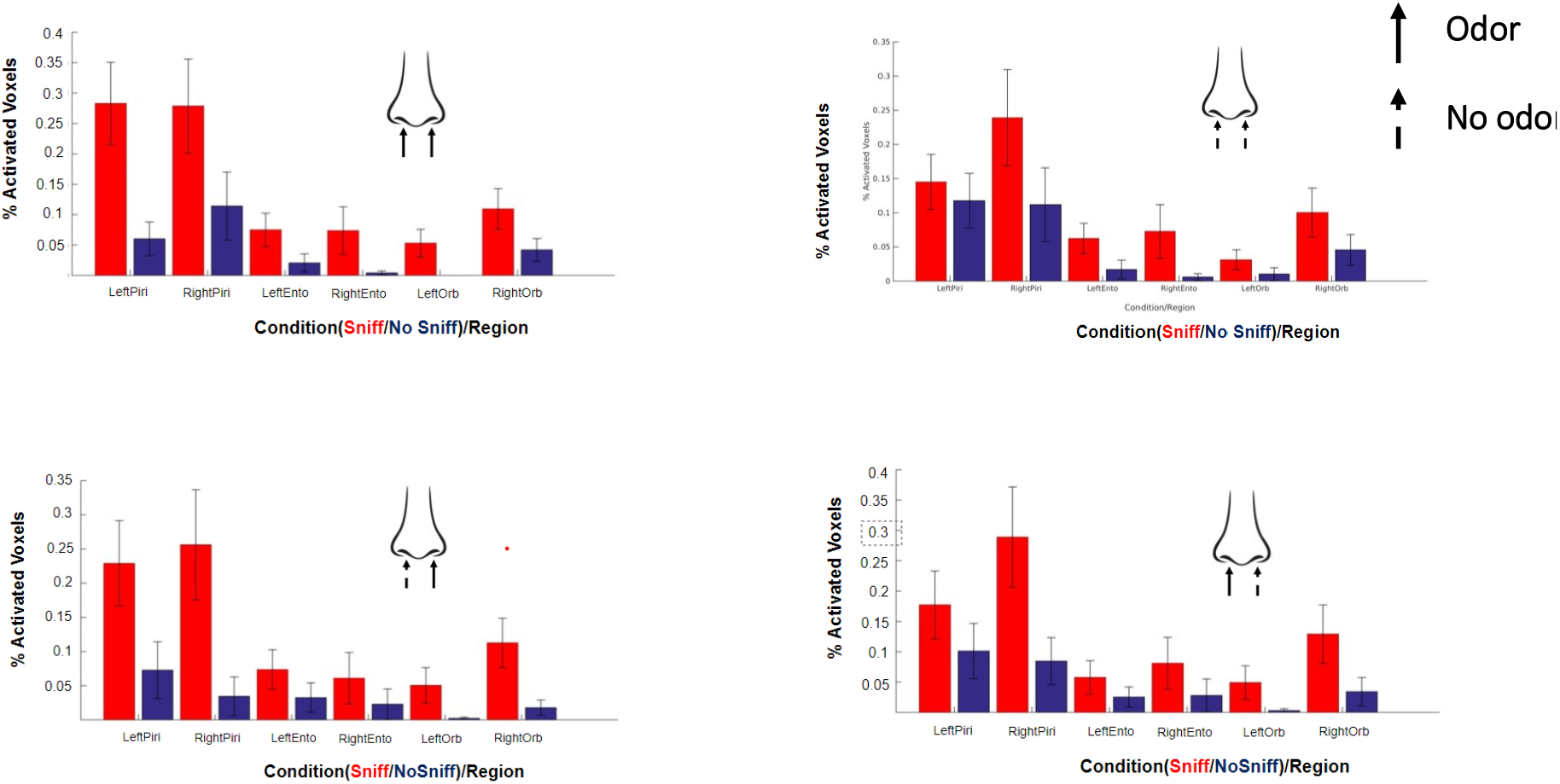
Percentage of activated voxels for each condition within the piriform, entorhinal and orbitofrontal cortices. Straight arrow denotes odor, dotted arrow denotes no odor. Red color denotes that subjects were sniffing and blue denotes that subjects were not sniffing.

When the frontal and temporal piriform activation was investigated using the ROIs shown in **Figure 3**, all three conditions elicited highly similar fMRI activation patterns. The contrasts between conditions yielded statistically insignificant differences even at very liberal statistical thresholds. Regardless of the stimulated nostril, fMRI activation in the temporal piriform cortex was bilateral (**Figure 6**, top row) with a larger cluster in the right hemisphere compared to the left. In contrast to the bilateral activation in the temporal piriform cortex, olfactory stimulation produced unilateral activation of the right frontal piriform cortex (**Figure 6**, middle row), and left entorhinal cortex (**Figure 6**, bottom row), each of which was consistent across all three stimulation conditions.

**Figure 6.**
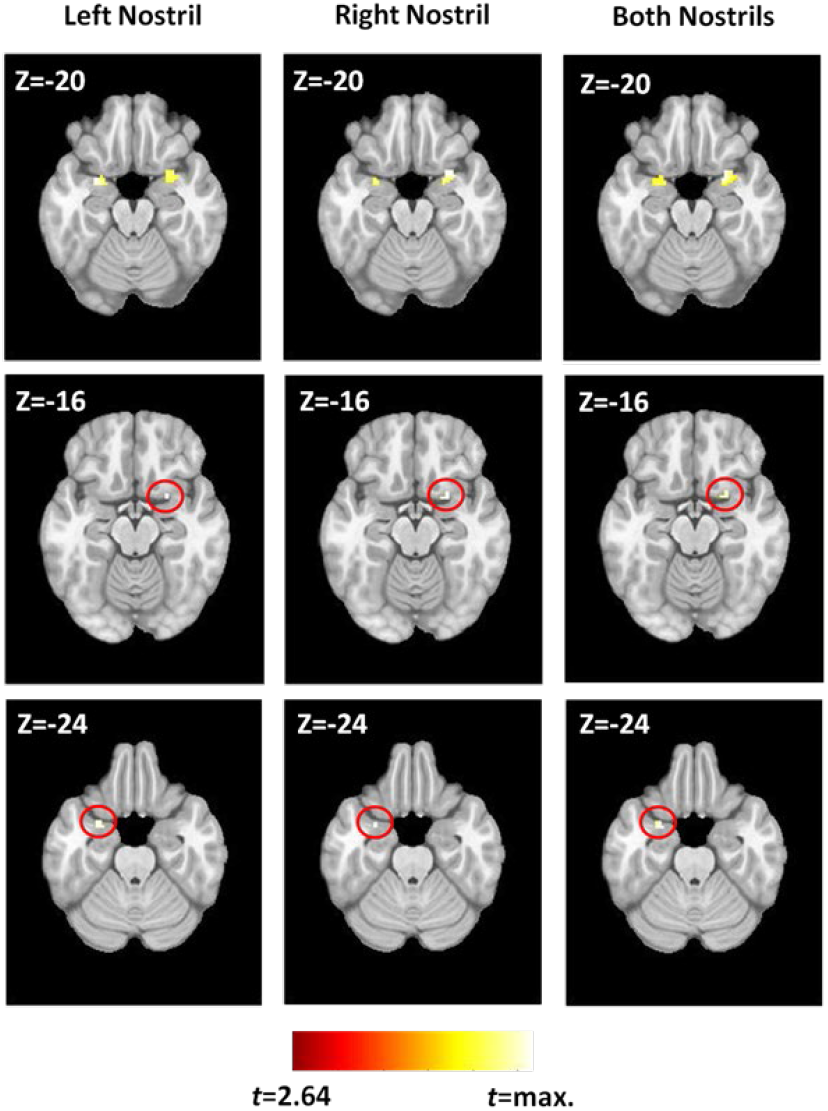
Activation in ROIs identified by the GLM.

Figure 7 shows the results of euSEM effective connectivity modeling with the four ROIs that were activated across all conditions. Only the contemporaneous stimulation effects are shown. The model indicates that the left temporal piriform influences both the right temporal piriform and the left entorhinal cortex. The right temporal piriform influences the right frontal piriform, which in turn influences the left entorhinal cortex. Finally, the left entorhinal cortex influences activity in the right frontal piriform cortex. As such, the right frontal piriform cortex has more inputs (including inputs from olfactory bulb, which are not shown) influencing its activation than the other network nodes, which may explain why it shows that largest volume of activated brain tissue.

**Figure 7.**
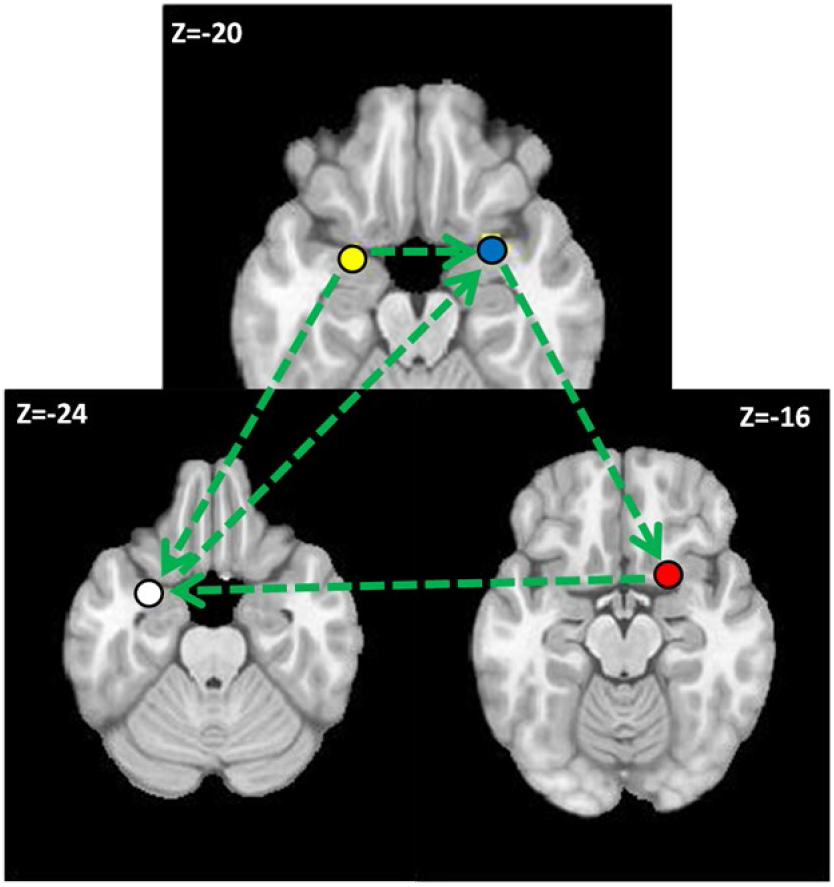
The effective connectivity model for odor processing.

## DISCUSSION

Based on olfactory system’s anatomical connectivity, it is reasonable to anticipate odor processing to be handled by the cerebral hemisphere that is ipsilateral to the stimulated nostril. However, the overall findings of this study do not support such a hypothesis and raise the possibility that secondary connections in the olfactory pathway may support contralateral odor processing. As mentioned in the introduction, there is evidence that the two olfactory tracts are linked through the anterior olfactory nuclei and the anterior commissure, which may allow contralateral odor processing to a certain extent (25).

In our experimental design, subjects simultaneously received clean air in one nostril and PEA in the other (26). While other odorants could also have worked, PEA has the benefit of being established as not localizable to a nostril (27). Therefore, we can assume that the fMRI tasks we used only involved very basic smell processing, such as detecting their presence or absence. This is different from complex tasks such as odor localization, which involves the intranasal trigeminal network as well.

In our data, there is little evidence to suggest that simply smelling a pure olfactory stimulus, whether presented bilaterally or unilaterally, produces asymmetrical neural activation patterns in the absence of cognitive challenges. When the no-sniff and sniff paradigms were contrasted, we can conclude that sniffing adds variability and amplitude to the fMRI signal during olfactory processing. Our connectivity analysis may explain how the primary and secondary anatomical connections in the olfactory system correspond to the functional pathways. For instance, the piriform cortex seems to be the primary olfactory processing region, and the entorhinal and orbitofrontal must be secondary olfactory regions. Activation of the piriform cortex (primary olfactory sensory region) may not necessarily activate secondary association regions unless it is necessary in the context of the current behavioral or environmental demands. As an alternative approach, future investigations could employ time-varying connectivity analysis to explore POC’s connectivity during nostril specific stimulation. It is important to note that the olfactory system employs reciprocal inhibition, depending on the type of stimulation (28, 29). Additionally, our study found that sniffing an odor or odorless air activates bilateral cerebellar activation, and similarly, smelling an odor or odorless air activates the anterior insula bilaterally, which may be related to salience monitoring.

Airflow changes stimulate the intranasal trigeminal system which is known to influence odor processing. Therefore, all of our experiments ensured that there were no differences in airflow rates between nostrils (via the use of flow meters). To minimize retronasal cross-contamination, future experiments can train study subjects to exhale from their mouths after inhaling odorants through their nose. For some participants in our study, we used T2-weighted images before and after fMRI scanning to evaluate and exclude subjects with nasal congestion and to ensure consistency of nostril dominance during experiments.

As per Wilson et al., the formation of odor objects is a result of experience-dependent changes mediated by plasticity changes in intracortical connectivity. These changes bind the activity of distributed cortical regions that correspond to a given olfactory stimulation (30). These divergent projections to olfactory cortical areas are able to transform odor information in many ways to form odor percepts. The olfactory cortex, therefore, is a crucial structure for translating inhaled odorants into rich emotion and memory tinged perceptions (30). For example, pleasant and unpleasant odorants are likely to impart changes in the olfactory system activity and connectivity configuration differentially. A null lateralized finding such as ours suggests that a given brain region, including its connections, may not exclusively support simple olfactory processing (31, 32). Therefore, future experiments should use different odorants and investigate how changes in hedonic, odor memory, or intensity ratings influence the olfactory system’s activity and connectivity patterns. Such experiments are critical to elucidate mechanisms by which the POC and other olfactory structures encode or process nostril specific olfactory stimulation.

According to Bar et al., the human brain can be considered as continuously generating predictions to approximate incoming sensory information (33). This is in contrast to passively ‘waiting’ to be activated by sensory information. Therefore, once sensory information of PEA is extracted from a nostril, the brain may link that information to stored memory representations. Such representations can subsequently activate associations relevant to PEA, resulting in a unique fMRI activity pattern irrespective of the stimulated nostril. This theory suggests that predictions help with perception and cognitive function by preparing relevant representations. It also explains why both primary and secondary olfactory structures are active bilaterally during monorhinal odor processing.

Clinically, the piriform cortex is considered a highly sensitive brain structure for the induction of epileptic seizures (34). Previously, this has been investigated in animal models by injecting certain pro-convulsant chemicals (35, 36). The piriform cortex may act as a secondary transfer point, transmitting epileptic discharges to other brain structures via the secondary olfactory pathway. For example, in the amygdala-kindling model of focal epilepsy, epileptic discharges from limbic areas to olfactory and non-olfactory regions (through the piriform cortex) can be observed (37). Therefore, it is important to investigate the contralateral connections of olfactory structures in order to understand how focal epilepsy becomes bilateral. The functional connectivity of olfactory structures identified in this fMRI study may be significant to understanding how epileptic discharges propagate contralaterally in the human brain.

In conclusion, the olfactory literature and our data analysis support the hypothesis that the perception of monorhinally presented odors is processed bilaterally in the brain, despite primarily having ipsilateral anatomical connections.

## ACKNOWLEDGEMENTS

This study was supported by NIH grants R01AG070088, R01NS099630, and R21AG064486.

## Notes

### Competing Interest Statement

The authors have declared no competing interest.

